# Application of Modular Response Analysis to Medium- to Large-Size Biological Systems

**DOI:** 10.1101/2021.07.27.453942

**Authors:** Meriem Mekedem, Patrice Ravel, Jacques Colinge

## Abstract

The development of high-throughput genomic technologies associated with recent genetic perturbation techniques such as short hairpin RNA (shRNA), gene trapping, or gene editing (CRISPR/Cas9) has made it possible to obtain large perturbation data sets. These data sets are invaluable sources of information regarding the function of genes, and they offer unique opportunities to reverse engineer gene regulatory networks in specific cell types. Modular response analysis (MRA) is a well-accepted mathematical modeling method that is precisely aimed at such network inference tasks, but its use has been limited to rather small biological systems so far. In this study, we show that MRA can be employed on large systems with almost 1,000 network components. In particular, we show that MRA performance surpasses general-purpose mutual information-based algorithms. Part of these competitive results was obtained by the application of a novel heuristic that pruned MRA-inferred interactions *a posteriori*. We also exploited a block structure in MRA linear algebra to parallelize large system resolutions.

**Author Summary:** The knowledge of gene and protein regulatory networks in specific cell types, including pathologic cells, is an important endeavor in the post-genomic era. A particular type of data obtained through the systematic perturbation of the actors of such networks enables the reconstruction of the latter and is becoming available at a large scale (networks comprised of almost 1,000 genes). In this work, we benchmark the performance of a classical methodology for such data called modular response analysis, which has been so far applied to networks of modest sizes. We also propose improvements to increase performance and to accelerate computations on large problems.

## Introduction

The expression and activity of genes and proteins in cells are controlled by highly complex regulatory networks involving genes and proteins themselves, but also non-coding RNAs, metabolites, etc. Despite tremendous efforts in research, including all the developments of high-throughput genomic technologies, a significant portion of this machinery remains uncharted. Moreover, dysregulations in such networks are related to many diseases, and healthy cells of a same organism feature adjusted regulatory networks depending on their cell types and states. Techniques, both experimental and computational methodologies, that enable the inference of regulatory networks for different cells are obviously of great interest.

Reference databases such as Reactome[1], KEGG[2], IntAct[3], or STRING[4] that compile our knowledge of biological pathways or protein interactions have been established that provide valuable reference maps. Due to their universal nature, these maps do not reflect natural and pathologic variations of regulatory networks though some chosen disease pathways might be included. In principle, researchers should generate data specific to the biological system of interest to assess the actual wiring of its regulatory network. Specific data can be combined with reference databases in some algorithms, while others only rely on *de novo* inferences. The field of systems biology has proposed many algorithms for such a purpose involving different modeling approaches[5–7]. Obviously, algorithms must match the type of data available to perform the inference such as a transcriptomes or proteomes obtained under multiple conditions, time series, or perturbation data.

In this work, we are interested in the inference of regulatory networks based on systematic perturbation data. That is, given a biological system of interest, which could be the whole cell, but also a small set of related genes or proteins such as a pathway or part of a pathway, we have access to information reporting the activity level of every component (gene/protein). Typical examples are transcript, protein, or phosphorylated protein abundances. This information is available in basal condition as well as under the systematic perturbation of each single component. When this type of data are obtained from a biological system in a steady state, modular response analysis[8] (MRA) has been widely and successfully applied[9]. The elegance of MRA is that it provides an efficient mathematical framework to estimate a directed and weighted network representing the system regulatory network. Most applications of MRA are limited to networks comprised of a modest number of modules (<10). In this study, we want to explore the application of MRA to medium-(>50) and large-size (>500) systems. It entails a particular implementation of the linear algebra at the heart of MRA to parallelize computations as well as the introduction of a heuristic to prune the inferred networks *a posteriori* to improve accuracy.

As stated above, rewiring of regulatory networks is natural and necessary to yield a multitude of cell types in higher organisms, and to adapt to distinct environmental conditions. Rewiring is also associated with several diseases[10,11], an extreme case being cancer[12–14]. For instance, kinase signaling cascades might be redirected in certain tumors to achieve drug resistance or to foster exaggerated cell growth. MRA has been applied to a number of such cancer-related investigations[15,16] considering rather small networks. Here, we take advantage of two published data sets that involve cancer cell lines and provide systematic perturbation data compatible with MRA requirements. The first – medium-size – data set[17] reports the transcriptional expression of 55 kinases and 6 non kinases under 11 experimental conditions (unstimulated plus 10 distinct stimulations). Under every condition, the transcript levels of all the 61 genes were obtained by shallow RNA sequencing, including wild type cells and cells with individual KOs of each gene. These data hence enable us to infer one network *per* condition (11 networks) to discover how those 61 genes regulate themselves transcriptionally. The second – large-size – data set was generated by the next generation of the Connectivity Map (CMap) using its new L1000 platform[18]. Both shRNA- and CRISPR/Cas9-based systematic perturbations of roughly 1,000, respectively 350, genes in 9, respectively 5, cell lines were released. These data enable us to infer 9+5=14 networks.

We compare the performance of MRA, with and without the proposed pruning heuristic, to mutual information (MI)-based methods that have found broad acceptance.

## Results

### Network inference algorithms

The availability of large functional genomics data collections (transcriptomes and/or proteomes) has led to the development of a number of algorithms aimed at inferring interaction networks [7]. An essential ingredient of most algorithms is the co-expression of genes (or proteins)[19], which can be captured by simple correlation coefficients[20], mutual information (MI), or diverse statistical models[21]. There are too many such algorithms to review them all here, but MI-based approaches seem to have provided off-the-shelf, robust solutions that are widely used. We hence compare MRA to representatives of this category such as CLR[22], MRNET[23], and ARACNE[24].

MI is often preferred over correlation for its ability to detect nonlinear relationships. With a network involving *n* genes whose expression levels are measured in *m* transcriptomes, we write *X*_*i*_ the discrete distribution representing gene *i* expression. The MI between genes *i* and *j* is given by

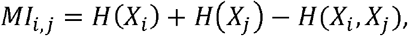

where *H*(*X*)= − ∑_*k* ∈ *X*_ *p* (*x*_*k*_) ln (*p* (*x*_*k*_)) is the entropy of a discrete random variable . There exist different estimators for *H*(*X*) that use the *m* available transcriptomes[25]. Networks of interactions identified though MI, imposing a minimal threshold on MI values, are commonly called relevance networks[26,27]. The CLR algorithm improves over relevance networks by introducing a row- and column-wise z-score-like transformation of *MI*_*i,j*_ to normalize the MI matrix into a *Z*= (*z*_*i,j*_)matrix before thresholding. Namely, for each gene *i* CLR computes

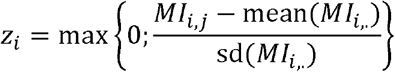

and then

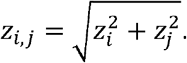

MRNET applies a greedy maximum relevance strategy to link each gene *i* to the gene *j* that has maximum MI with it (*j* = arg max*MI*_*i,j*_). Additional links are added recursively maximizing MI with both the gene *i* and the already linked genes until a stop criterion based on redundancy is met. A further approach by pruning was proposed by ARACNE authors, where as in relevance networks a common threshold is applied to all the *M*_*i,j*_ followed by the application of a pruning rule. This rule states that, if gene *i* interacts with gene *j* through gene *k*, then, *M*_*i,j*_ ≤ min{*M*_*i,k*_; *M*_*k,j*_}. Consequently, among each triplet of nonzero MI after initial thresholding, the weakest interaction is removed.

### The MRA and MRA+CLR algorithms

Due to its ability to model biological systems at various resolutions, the MRA terminology for a system component is a module. We follow this terminology and consider that the *n* modules composing the system have their activity levels denoted by *x* ∈ ℝ^*n*^. Here, modules are genes and *x*_*i*_ stands for gene *i* transcript abundance. If we make the rather nonrestrictive assumption that relationships between modules are modeled by a dynamical system

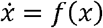

(*f* (.)must exist but it does need to be known), and the system is in a steady state at the time of experimental measurements 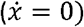, MRA theory lets us compute an *n* × *n* matrix of local interaction strengths *r*=(*r*_*i,j*_) from a gene *j* to a gene 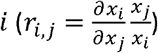. The matrix *r* is obtained from linear algebraic computations based on the observed activity of each module in an unperturbed state, and under the individual, successive perturbations of each module. Details are provided in MRA original publication[8], reviews of MRA developments[9], or in our recent publication[15]. We use the notations of this recent paper. In Materials and Methods, we provide a brief overview of MRA along with a description of the particular way we implemented the linear algebra to take advantage of parallel computing.

Returning to the regulatory network inference problem, the MRA local interaction matrix *r* provides us with a direct estimate of this network. Interactions are signed with positive coefficients representing activation and negative coefficients representing inhibition. Given the fact that we want to apply MRA to large systems, where every module does not necessarily have a direct influence on all the others, we also face the problem of thresholding or pruning. Within the context of this study, we call MRA the direct use of MRA computations followed by a threshold on the absolute values of *r* coefficients (values below a given threshold in absolute values are set to 0). We also adapted CLR heuristic (z-score-like computation) to bring *r* coefficients to a more uniform scale before thresholding. We call this algorithm MRA+CLR, see Materials and Methods for details.

### Application to a medium-size data set

Gapp *et al*.[17] published a data set, where they studied the transcriptional impact of the full knockouts (KOs) of 55 tyrosine kinases and 6 non-kinases. We call this data set K61. The systematic perturbations (KOs) of each gene as well as the unperturbed transcriptomes obviously constitute a *bona fide* MRA data set. The transcriptomes were acquired under 11 conditions: no stimulation (None), FGF1, ACTA, BMP2, IFNb, IFNg, WNT3A, ionomycin (IONM), resveratrol (RESV), rotenone (ROTN), and deferoxamine (DFOM) stimulation. Stimulations were applied for 6 hours allowing the cells to adapt and reach a steady state or near steady state. To facilitate the generation of full-KOs, human HAP1 haploid cells[28] were utilized. The published transcriptomes were not limited to the expression of the 61 perturbed genes, but here, due to the specifics of MRA, we limited the data to those 61 genes. Replicates were essentially averaged (see Materials and Methods), resulting in a 61 × 61 matrix for each of the 11 conditions. Interestingly, considering the complete transcriptomes, K61 authors showed in their publication that those clustered primarily after the stimulatory condition. That is transcriptomes of different KOs obtained under the same stimulation were closer to each other than transcriptomes of the same KO but under different conditions. When reduced to the 61 genes of the network, this picture was less pronounced. In Fig. 1, we see that None-, WNT3A-, and to a certain extent IFNg-stimulated transcriptomes clustered separately thus potentially indicating rather different network wiring. The other conditions were not really separated suggesting that more similar networks could take place.

**Figure 1.**
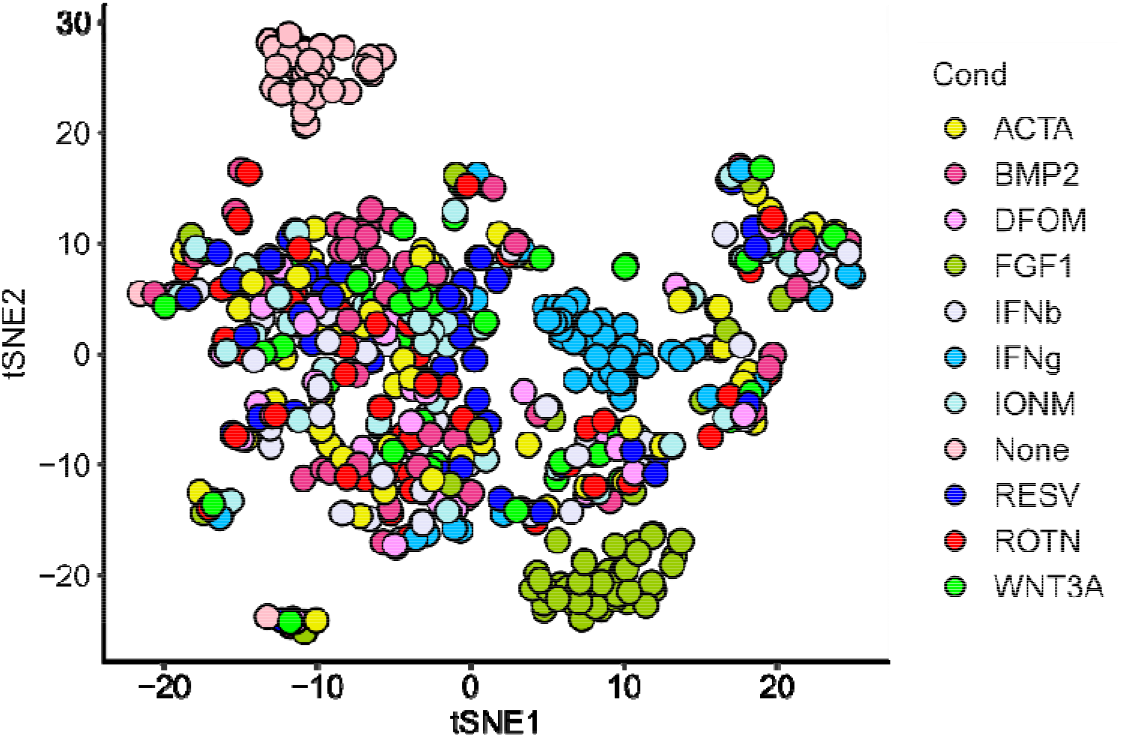
t-distributed stochastic neighbor embedding (t-SNE) 2D projection of the 61 11 transcriptomes of the K61 data set.

We applied MRA, MRA+CLR, CLR, MRNET, and ARACNE to the K61 data set, the later 3 algorithms implementations were provided by the minet BioConductor package[25]. To estimate performance, we compared our results with the STRING database[4] due to its broad content. In fact, working with transcriptomic data, the inferred networks might overlap protein complexes as well as certain parts of known pathways, but they might also unravel different types of relationships such as genetic interactions, strong co-regulation, etc. Physical interaction of well-described pathway databases[1,3] might thus be too restrictive. To apply a uniform selection mechanism to all of the algorithms, we simply took the top 5%, 10%, 20%, 30% and 40% scores of the returned interaction matrices and determined the intersection with STRING. This resulted in confusion matrices reporting true/false positives (TPs/FPs) and true/false negatives (TNs/FNs) along with a P-value for the significance of the STRING intersection (hypergeometric test). A representative example (None condition) is featured in Fig. 2A, while the complete results are in Suppl. Table 1. Given the limited overlap between STRING and our data, and the rather large numbers involved in the confusion matrices, we found the P-values rather unstable (small differences in confusion matrices might cause important changes in terms of P-values). They should hence be regarded as indicative only. Because we used a constant reference (STRING), and all the algorithm scores were selected in identical numbers, reporting the number of TPs gives a clear indication of the relative algorithm performances. In Fig. 2B-E we provide these numbers at the top 10% and the top 20% selection levels. ARACNE implementation in minet did not perform well, typically reaching half of CLR or MRNET TPs. Accordingly, ARACNE performance is not reported in Fig. 2, but in Suppl. Table 1 only. The CLR heuristic applied on top of MRA did not provide much performance increase, but it resulted in more stable performances thus making it nonetheless an attractive option.

**Figure 2.**
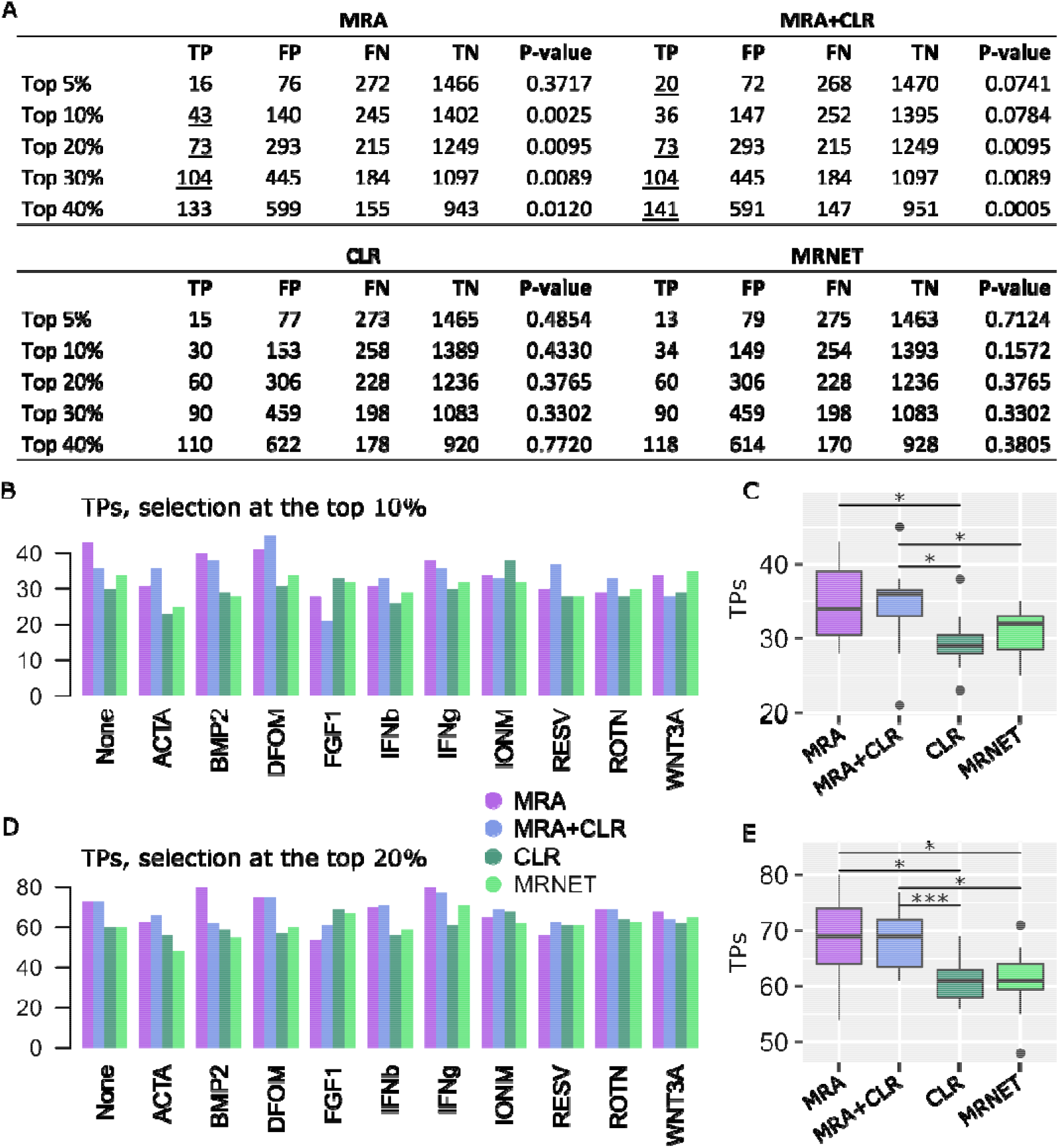
Performance on K61 data. **A**. Representative confusion matrices for the None condition. **B**. TP numbers at the top 10% selection level. **C**. Comparison between the algorithm TP numbers (Wilcoxon test, 2-sided, *P < 0.05). **D**. TP numbers at the top 20% selection level. **E**. Comparison between the algorithm TP numbers (Wilcoxon test, 2-sided, *P < 0.05, ***P<0.005).

In their article, K61 authors discussed interesting differences in JAK1 *versus* JAK2 and TYK2 signaling, three members of the JAK family. In particular, they found that JAK1 KO cells were insensitive to IFNb and IFNg stimulation, while JAK2 and TYR2 KO cells responded normally although, in general, all these proteins are known to contribute to transcriptional response upon type I and II interferon stimuli[29]. To illustrate how network inference might provide some clue on such differences, we report in Fig. 3A the MRA+CLR-inferred transcriptional interaction strengths between those three genes and their targets under the unstimulated (None), IFNb, and IFNg conditions. In the absence of stimulation, we clearly notice opposed influences of JAK1 on its targets compared to JAK2 and TYR2 (first three columns), which already indicate different signal transduction capabilities. Upon IFNb stimulation, the interactions are closer with opposed action on ROR1 and PDFGRA. JAK2 and TYR2 remained highly similar in this condition. IFNg stimulation induced three different patterns with ROR1 transcriptional inhibition remaining a specific mark of JAK1. Gapp *et al*. also found differences in FGF receptors. FGF-induced response was attenuated in FGFR1 and FGFR3 KO cells, but preserved in FGFR2 and FGFR4 KO cells. In Figure 3B, we notice an almost perfect inversion of the activation/inhibition pattern between FGFR1 *versus* FGFR2 and FGFR3. FGFR4 adopted a very different configuration with limited interactions. This observation already indicates a distinct role for FGFR1. Upon FGF stimulation, the interactions are more patchy, but certain oppositions can be found such as a strong inhibitory action of FGFR1 and FGFR3 on RYK transcription.

**Figure 3.**
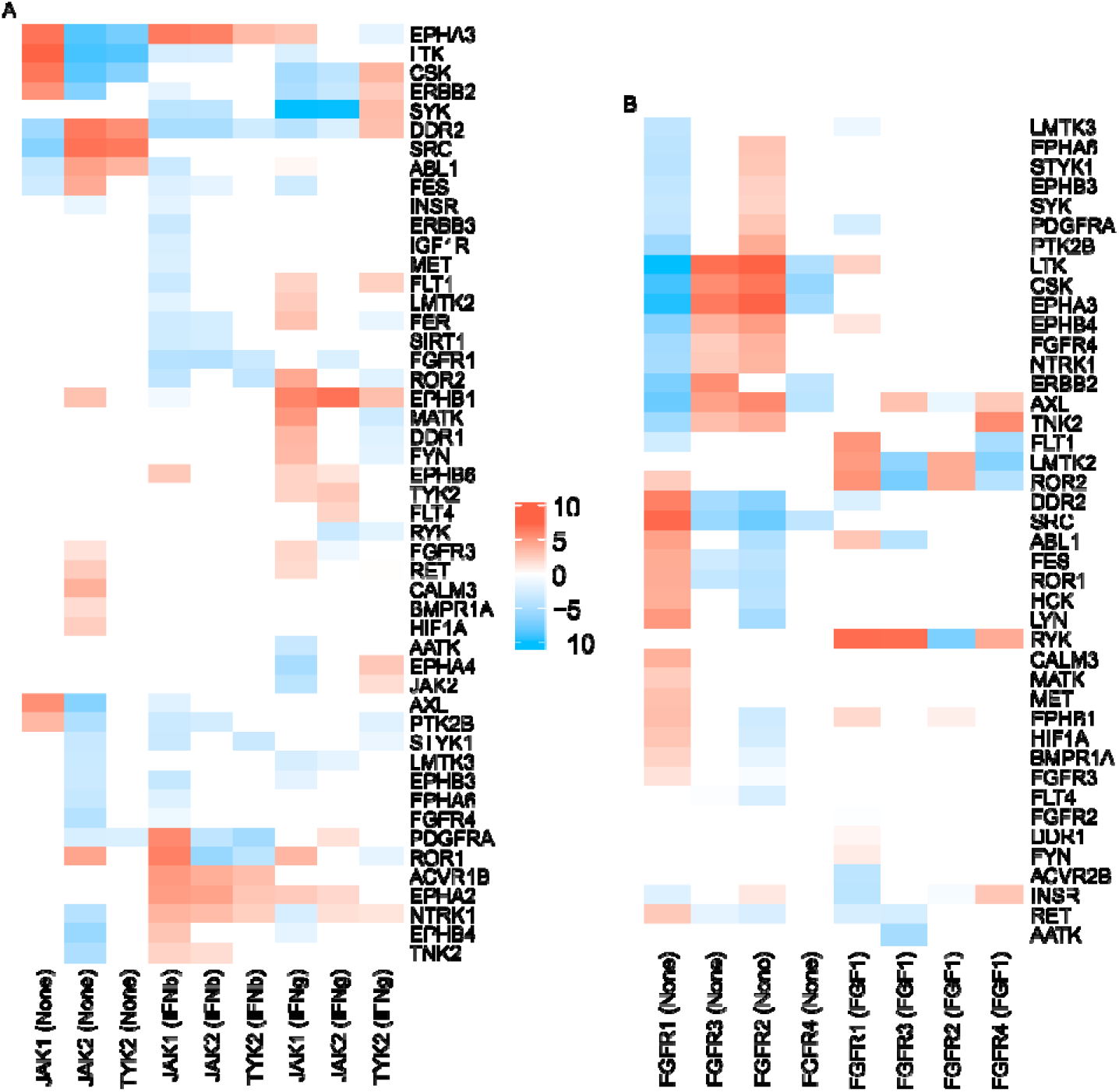
MRA+CLR-inferred interactions (top 20% selected). **A**. Interaction strengths (in log_2_ with sign preserved) between JAK1, JAK2, and TYR2 and their targets. Stimulatory conditions are in brackets (None, IFNb, IFNg) **B**. Interaction strengths between FGFR1, FGFR2, FGFR3, and FGFR4 and their targets.

### Application to a large-size data set

CMap next generation platform L1000[18] has recently released (December 2020) a new batch of data. These data are in majority comprised of transcriptomes obtained in reference cancer cell lines under a large number of perturbations with chemical agents, but most importantly shRNA-induced knockdowns and CRISPR/Cas9 KOs. L1000 cost effective design entailed the identification of roughly 1,000 *hallmark* genes from which a large proportion of the whole transcriptome can be inferred. The L1000 platform only measures the expression of the hallmark genes experimentally. Two subsets of these data interest us.

A first data set is composed of the almost systematic shRNA perturbation of all the hallmark genes, thus providing an expression matrix close to 1,000 ×1,000 in size for 9 human cell lines: A375 (metastatic melanoma), A549 (lung adenocarcinoma), HCC515 (non-small cell lung cancer, adenocarcinoma), HT29 (colorectal adenocarcinoma), HEPG2 (hepatocellular carcinoma), MCF7 (breast adenocarcinoma), PC3 (metastatic prostate adenocarcinoma), VCAP (metastatic prostate cancer), and HA1E (normal kidney cells). To alleviate shRNA off-target effects, L1000 employed multiple hairpins, which were integrated into a consensus gene signature (CSG) that the authors showed to be essentially devoid of off-target consequences[18]. Cells were harvested 96 hours after shRNA perturbation leaving time to reach a steady state that is compatible to shRNA common use. Due to variation in data production, the actual matrix sizes ranged from 815 ×815 (MCF7) to 938 ×938 (A375). Interestingly, the t-SNE 2D projection of all the L1000 shRNA transcriptomes used here clearly indicate cell line specific subnetworks as well as shared, core parts (Fig. 4).

**Figure 4.**
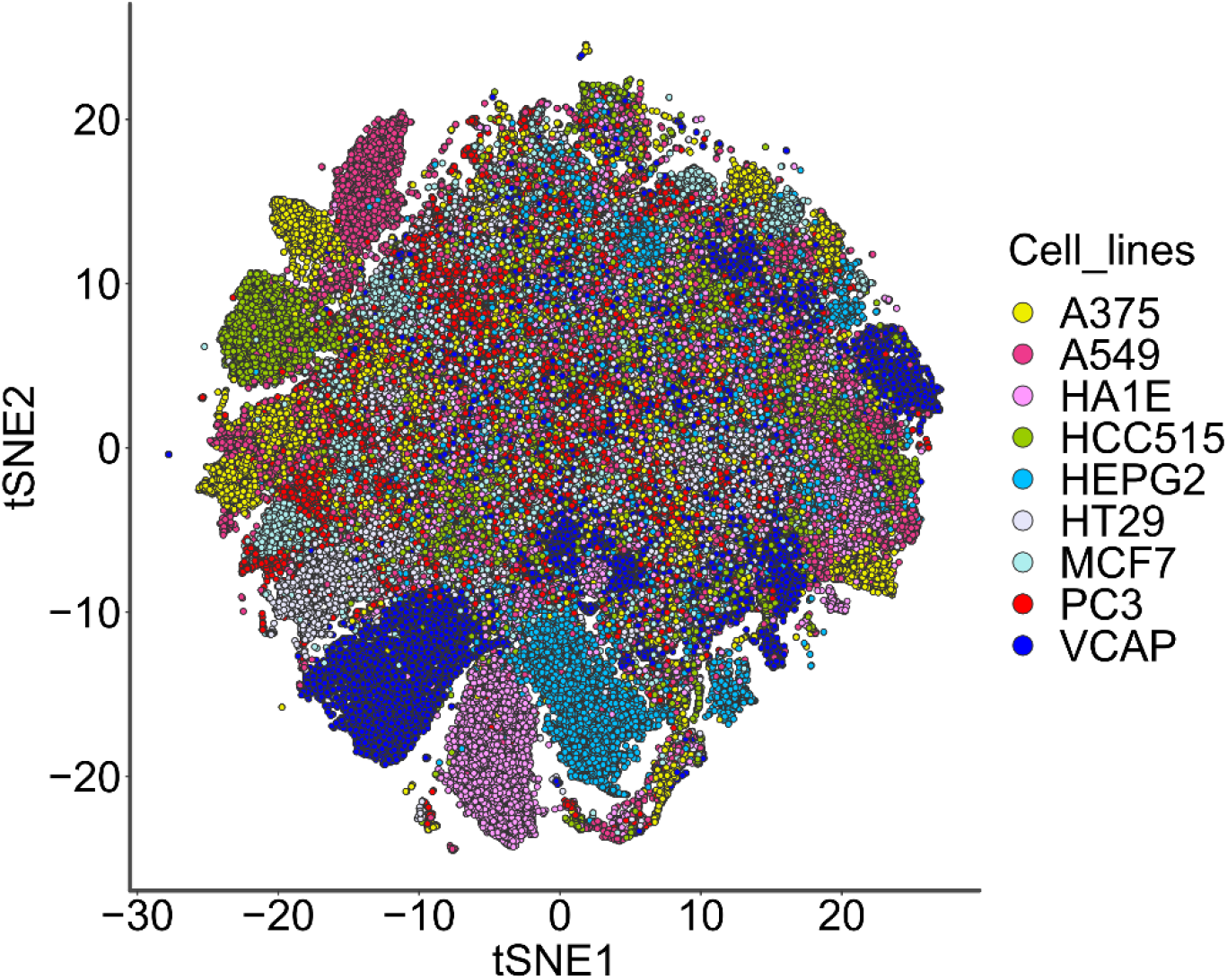
t-SNE projection of L1000 shRNA data. We note well-separated clusters that are specific to certain cell lines, *e*.*g*., HA1E, VCAP, HCC515, HEPG2, A549, A375, as well as shared undistinguishable profile. This indicates potential common and specific subnetworks across the cell lines.

We followed the same performance evaluation procedure as above for K61. A representative (A375 cells) confusion matrix is reported in Fig. 5A (full results in Suppl. Table 2), followed by TP numbers at the top 10% and top 20% selection levels in Fig. 5B-E. With these larger matrices, but also knockdown perturbations instead of KOs, MRA and MRA+CLR advantage was much augmented. Moreover, the CLR heuristic not only attenuated performance variability, but it almost systematically outperformed MRA alone.

**Figure 5.**
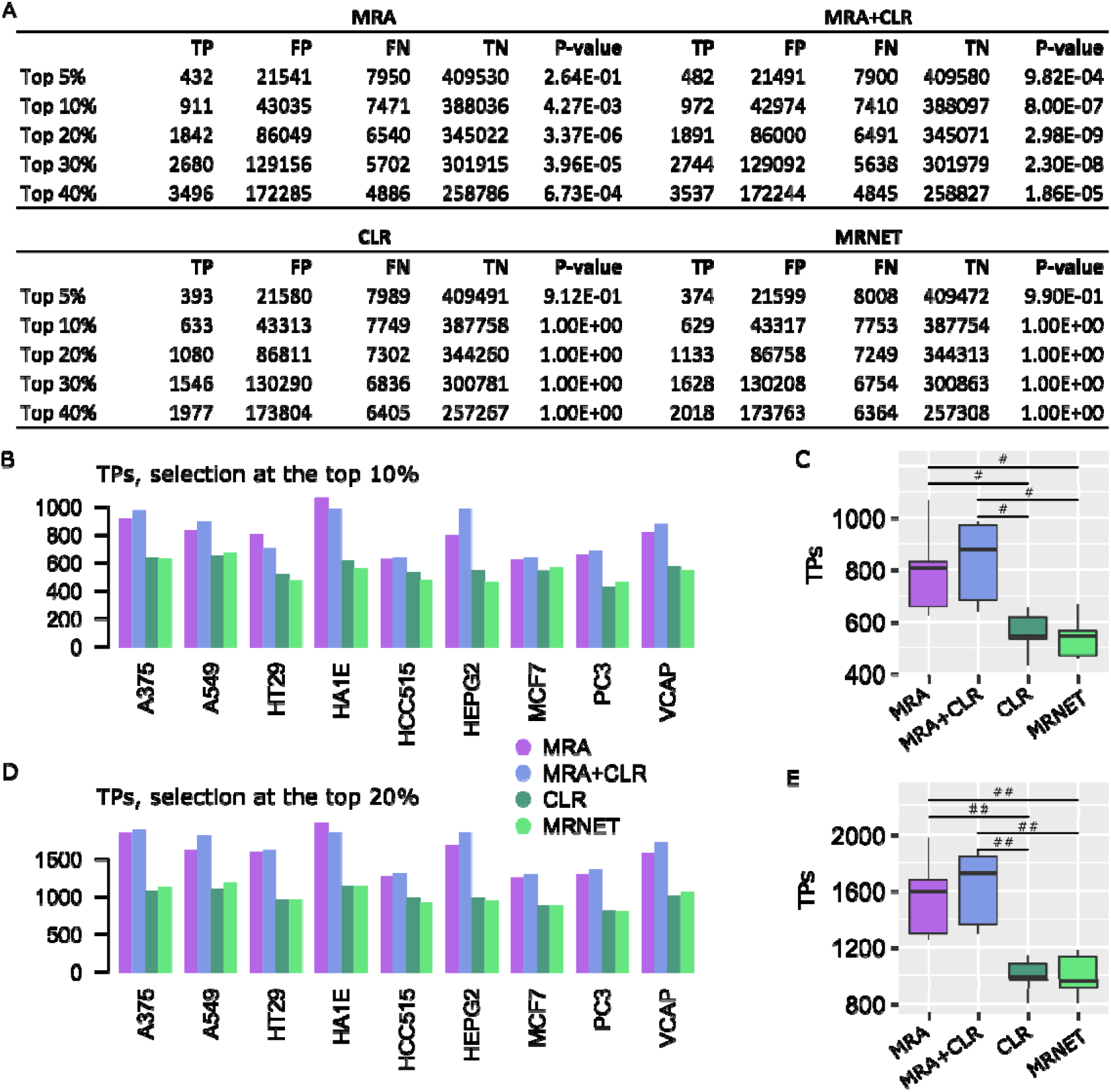
Performance on L1000 shRNA data. **A**. Representative confusion matrices for A375 cells. **B**. TP numbers at the top 10% selection level. **C**. Comparison between the algorithm TP numbers (Wilcoxon test, 2-sided, #P < 0.001). **D**. TP numbers at the top 20% selection level. **E**. Comparison between the algorithm TP numbers (Wilcoxon test, 2-sided, #P < 0.001, ##P < 0.00005).

To illustrate the interest of network inference at this scale, we intersected MRA+CLR inferences in normal kidney HA1E and melanoma A375 cells with a Gene Ontology term, *i*.*e*., GO:0006974 cellular response to DNA damage stimulus. In Fig. 6, we can notice the difference in connectivity between normal cells and cells where this process is obviously exacerbated, in particular the regulation of ATMIN a key molecule in DNA repair. This result is in agreement with the known rewiring of genetic networks in response to DNA damage[30].

**Figure 6.**
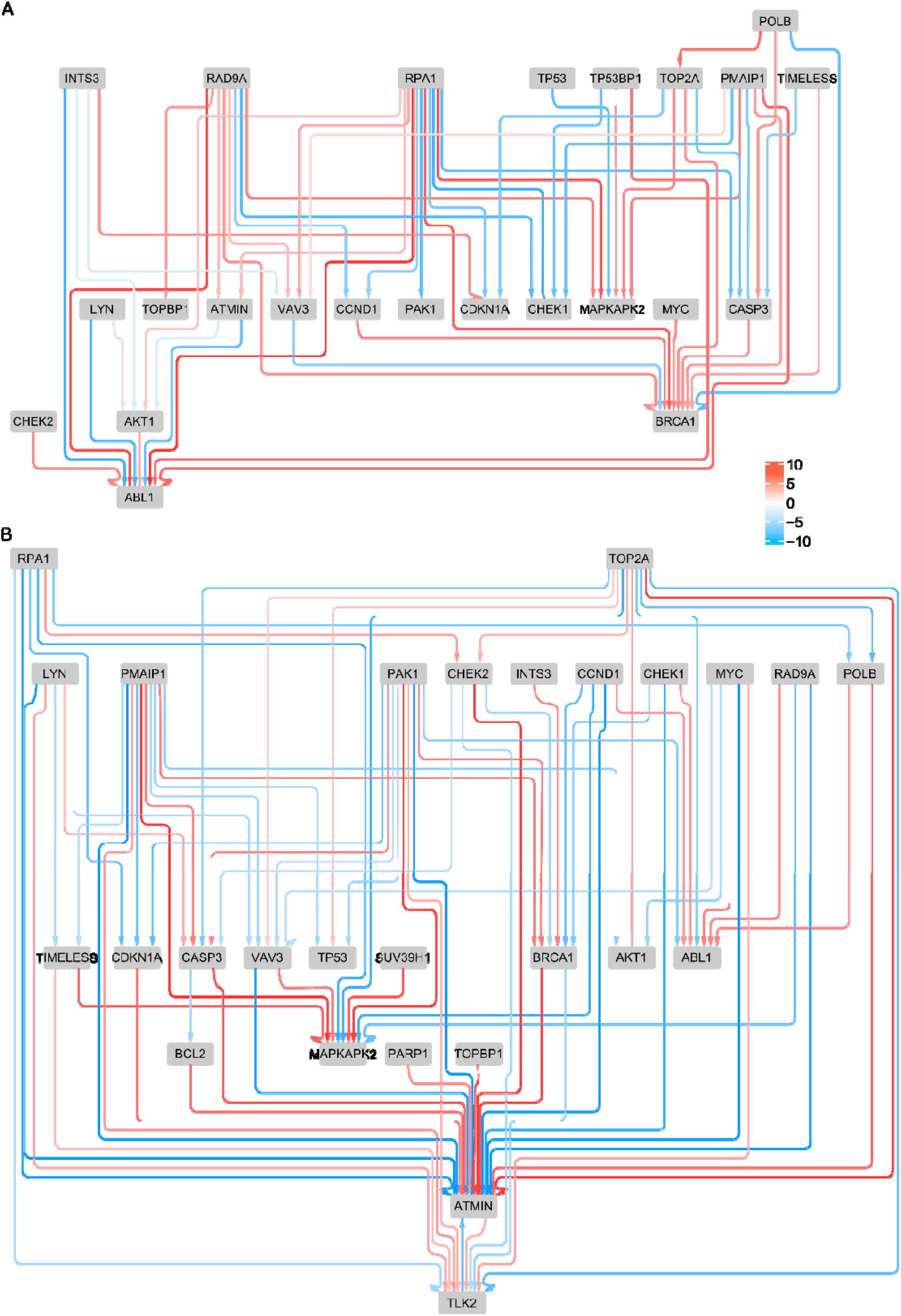
Networks inferred with MRA+CLR (top 10% selection) in normal kidney cells (**A**) and melanoma cells (**B**) for genes involved in cellular response to DNA damage stimulus (GO:0006974).

The second L1000 data set of interest is the CRISPR/Cas9 collection of KOs. These data were only available for five cell lines: A375, A549, HT29, MCF7, and PC3. The matrix sixes ranged from 343 343 (MCF7) to 359 359 (A375). Performance results are featured in Fig. 7 and Suppl. Table 3. Although MRA and MRA+CLR again dominated the other algorithms, their advantage was less pronounced on these large, full KO data.

**Figure 7.**
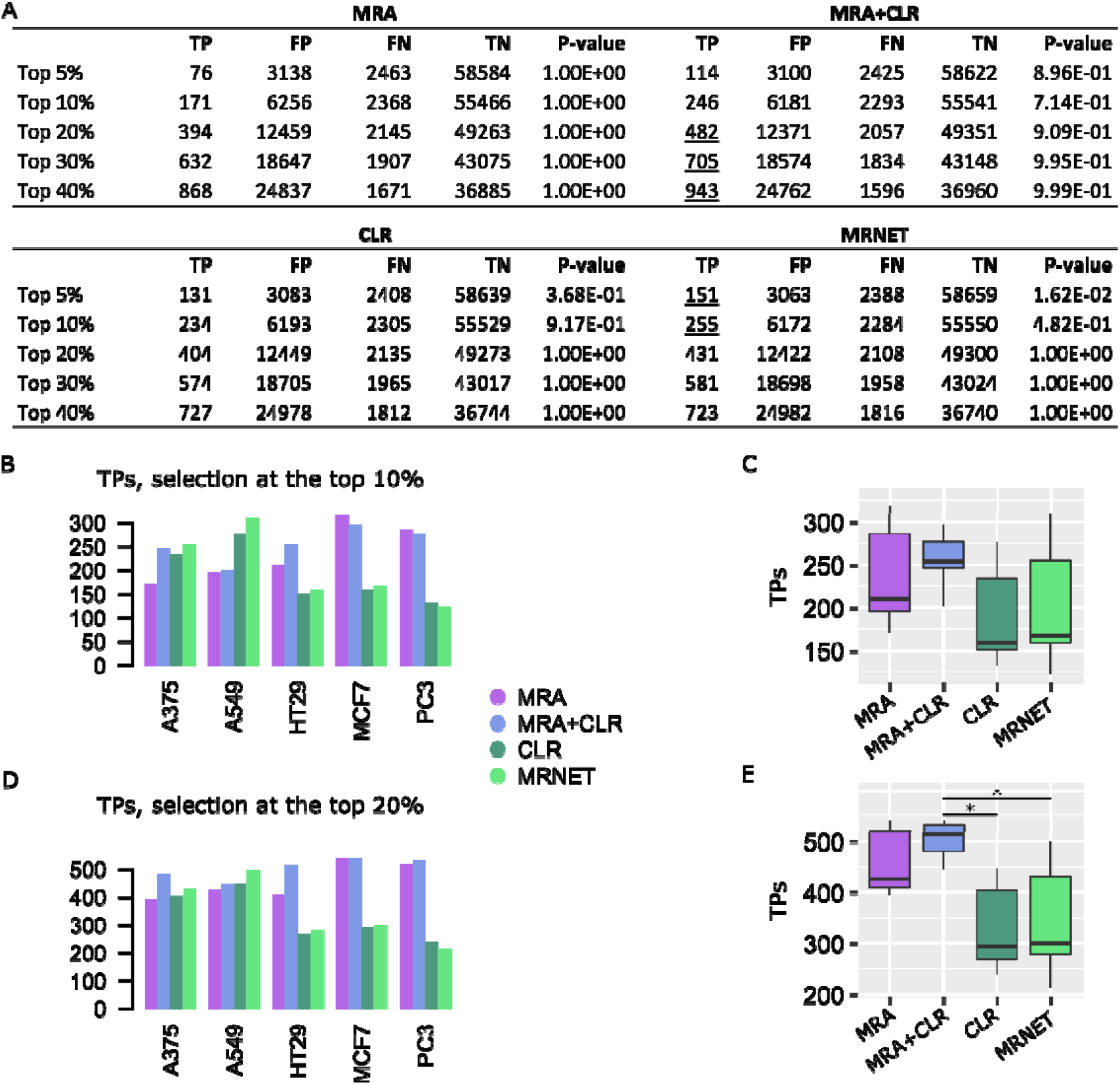
Performance on L1000 CRISPR/Cas9 data. **A**. Representative confusion matrices for A375 cells. **B**. TP numbers at the top 10% selection level. **C**. Comparison between the algorithm TP numbers. **D**. TP numbers at the top 20% selection level. **E**. Comparison between the algorithm TP numbers (Wilcoxon test, 2-sided, *P < 0.05).

## Discussion

We presented a particular application of MRA to large biological systems and showed its competitive performance compared to first-in-class MI-based inference methods. Obviously, MI-based methods have a much broader spectrum of application, as they do not need specific and systematic perturbations on the components of the biological system whose network is inferred. Nevertheless, when perturbation data are available, our results suggest that a dedicated method, relying on a modeling approach might deliver good performance in a robust fashion. The simple heuristic we proposed to prune MRA inferences, which was adapted from the CLR algorithm, provided more stability in MRA performance. In many cases, especially with very large systems (*n ≈* 1,000), this heuristic boosted performance.

Although the number of data sets was limited, we could notice much superior improvement over MI-based methods with L1000 shRNA knockdown perturbation data compared to the two full KO data sets. This might relate to the linearization at the heart of MRA modeling, where the error depends on the magnitude of perturbations (see our derivation of MRA through Taylor series expansion[15]). Very strong perturbation such as full KOs might bring the data away from MRA area of safe application.

## Materials and Methods

### Modular response analysis

We briefly recall the main MRA equations to facilitate the reading of this text, and to explain the particular way we implemented the linear algebra. We assume that the biological system is comprised of *n* modules whose activity levels are denoted by *x* ∈ℝ ^*n*^. We further admit the existence of *n* intrinsic parameters, *p* ∈ℝ^*n*^, one per module, and each of them can be perturbed by an elementary perturbation. One can imagine *x* reporting mRNA abundances and perturbations induced by shRNAs for instance. Lastly, we assume that there exist *S* ⊂ ℝ ^*n*^ × ℝ ^*n*^, an open subset, and *f*:*S* → ℝ ^*n*^ of class 𝒞 ^1^,*i*.*e*., continuously differentiable, such that

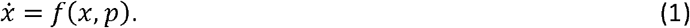

We do not need to know *f*(*x,p*) = (*f*_1_(*x,p*),…,*f*_*n*_(*x,p*))^*t*^ explicitly, but we need the existence of a time *T*> 0 such that all the solutions, for any *p* and initial conditions of *x*, have reached a steady state, *i*.*e*.,

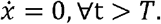

The unperturbed, basal state of the modules is denoted *x*(*p*^0^) ∈ ℝ^*n*^and it has corresponding parameters *p*^0^ ∈ ℝ^*n*^. By the application of the implicit function theorem and Taylor expansion at the first order [8,15], MRA relates the experimental observations of the global effect of perturbations to local interaction strengths, *i*.*e*., the matrix 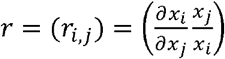 that we mentioned in Results. Such local interactions are obviously signed and non-symmetric. To compute *r*, we need to compute the relative global change induced by each elementary perturbation in each module. These values are compiled in a *n × n*matrix denoted *R* = (*R*_*i,j*_)with

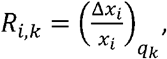

the relative difference in activity of module upon Δ*p*_*k*_ change induced by an elementary perturbation *q*_*k*_ that touches module *k* only. The relationship between observational data in *R* and the local interactions we want to estimate in *r* are provided by the following equations

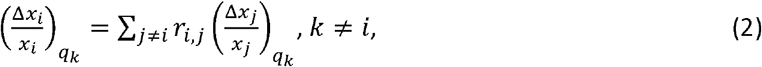

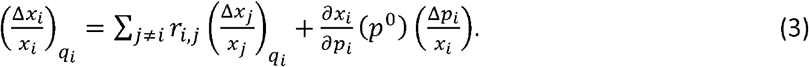

By setting *r*_*i,i*_ = −1, Eqs (2) and (3) can be put together in matrix form and we obtain

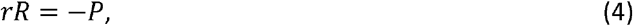

where *P* is a diagonal *n× n* matrix with

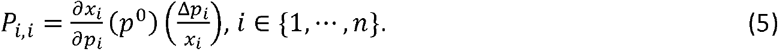

Eq. (3) can be solved in two steps: *r*= −*PR*^−1^and *r*_*i,i*_ = −1 imply *P*_*i,i*_ (R^−1^)_*i,i*_ = 1, thus

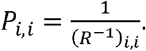

Therefore,

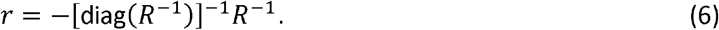

In practice, relative differences in *R* are often estimated with the more stable formula

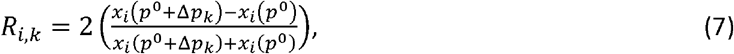

where we denote *x*(*p*^0^+Δ*p*)the steady-state corresponding to the changed parameters *p*^0^+Δ*p, i*.*e*., the solution of 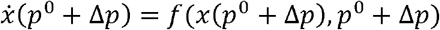.

### Parallelized and stable linear algebra

Eq. (6) requires the computation of the inverse of the matrix *R*, which is less efficient and less stable than LU decomposition with pivot search[31]. These technical issues are usually irrelevant with small systems, but in applications of MRA to larger biological systems they should be addressed.

As several authors noticed, including in MRA original publication[8], the homogeneous Eq. (2) is sufficient to compute *r*. Moreover, letting take the values1,…, *n*, we remark that Eq. (2) defines *n* systems of linear equations of dimension *n* −1, which can be solved independently. In particular, those systems can be solved on independent processors by performing the LU decomposition with pivot search. Illustrative speedup curves are featured in Fig. 8. Depending on the size of *n*, each such subsystem could itself benefit from a parallel solver if enough processors were available.

**Figure 8.**
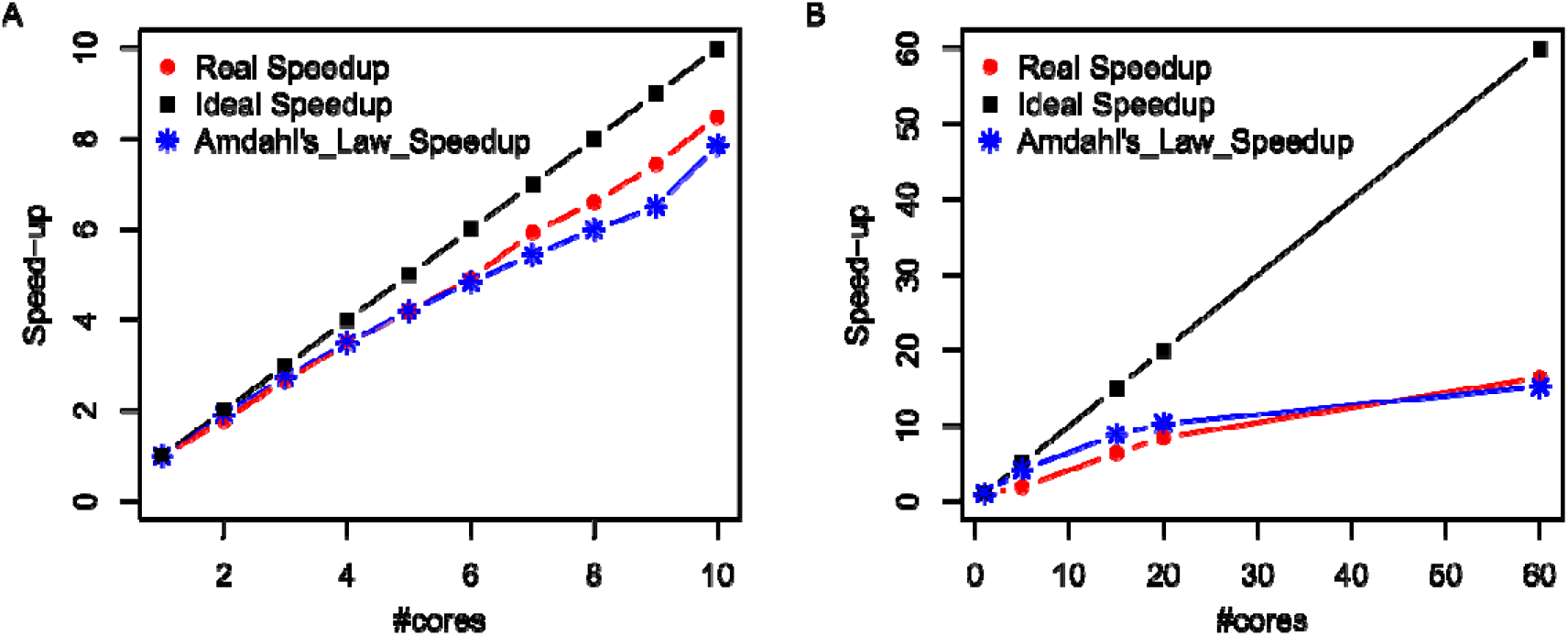
Speedup curves. **A**. K61 data(None condition, 61 61 matrix). **B**. L1000 shRNA data (A375 cells, 938 938 matrix).

When Eq. (2) is solved for each value of *i*, it is straightforward to solve Eq. (3) to find *P*_*i,i*_ values in case those are required:

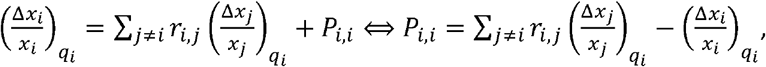

where Eq. (4) was used for the definition of .

### CLR, MRNET, and ARACNE computations

We used the implementation of these algorithms provided by the BioConductor R package minet[25]. The performance reported here reflects the performance of this specific implementation.

### CLR heuristic adapted to MRA

We adapted the CLR normalization scheme by means of z-score computation to MRA matrix content. From we thus derive a defined as follow:

_, with the standard deviation of ’s -th row,

_, with the standard deviation of ’s -th column,

_,and

### Data sets preparation

TK61 data were obtained on multiple 96-well plates. Accordingly, we tried to stick to this format preparing data for MRA computations. We computed an *R* matrix for each plate and then simply averaged the relevant *R*’s for each experimental condition to obtain the averaged *R* used in MRA. For MI-based inferences, we averaged all the relevant values.

L1000 shRNA data were extracted at level 5 (L1000 terminology) where CGSs (integration of multiple shRNA hairpins to alleviate off-target effects) were transformed into z-scores for normalization purposes by the authors of the data. Consequently, values representing the abundance of a gene were no longer positive numbers but just real numbers. Eq. (7) above was adapted to compute the relative changes in MRA *R* matrices according to

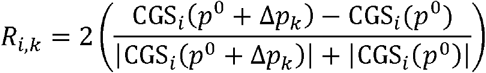

avoiding potential divisions by 0 in case of small values with opposed signs.

L1000 CRISPR/Cas9 data were averaged over replicates (also level 5).

### Performance evaluation

STRING as well as MI-based inference are devoid of direction of interaction and a sign. Therefore, the intersection of inferences with STRING content only used the upper triangular part of matrices representing the inferences (such matrices are symmetric anyway). To provide a fair comparison with MRA and MRA+CLR, we filled the upper triangular part of *r* according to *r*_*i,j*_.= max{| *r*_*i,j*_ |; | *r*_*i,j*_ |}, *i < j*

## Acknowledgements

MM was supported by a PhD fellowship of the Algerian government.

## Data availability

Data used in this work were made publicly available by their respective authors.

## Supporting information caption

**Supplementary Table 1**. Confusion matrices on the K61 data set.

**Supplementary Table 2**. Confusion matrices on the L1000 shRNA data set.

**Supplementary Table 3**. Confusion matrices on the L1000 CRISPR/Cas9 data set.

## References

1. Matthews L, Gopinath G, Gillespie M, Caudy M, Croft D, de Bono B, et al. Reactome knowledgebase of human biological pathways and processes. Nucleic Acids Res. 2009;37: D619–22. doi:10.1093/nar/gkn863

2. Kanehisa M, Araki M, Goto S, Hattori M, Hirakawa M, Itoh M, et al. KEGG for linking genomes to life and the environment. Nucleic Acids Res. 2008;36: D480–4.

3. Kerrien S, Alam-Faruque Y, Aranda B, Bancarz I, Bridge A, Derow C, et al. IntAct--open source resource for molecular interaction data. Nucleic Acids Res. 2007;35: D561–5. doi:10.1093/nar/gkl958

4. Szklarczyk D, Franceschini A, Kuhn M, Simonovic M, Roth A, Minguez P, et al. The STRING database in 2011: functional interaction networks of proteins, globally integrated and scored. Nucleic Acids Res. 2011;39: D561–8. doi:10.1093/nar/gkq973

5. Bansal M, Belcastro V, Ambesi-Impiombato A, di Bernardo D. How to infer gene networks from expression profiles. Molecular Systems Biology. 2007;3: 78. doi:10.1038/msb4100120

6. Babtie AC, Stumpf MPH, Thorne T. Gene Regulatory Network Inference. In: Wolkenhauer O, editor. Systems Medicine. Oxford: Academic Press; 2021. pp. 86–95. doi:10.1016/B978-0-12-801238-3.11346-7

7. Emmert-Streib F, Dehmer M, Haibe-Kains B. Gene regulatory networks and their applications: understanding biological and medical problems in terms of networks. Front Cell Dev Biol. 2014;2. doi:10.3389/fcell.2014.00038

8. Kholodenko BN, Kiyatkin A, Bruggeman FJ, Sontag E, Westerhoff HV, Hoek JB. Untangling the wires: a strategy to trace functional interactions in signaling and gene networks. Proc Natl Acad Sci U S A. 2002;99: 12841–6. doi:10.1073/pnas.192442699

9. Santra T, Rukhlenko O, Zhernovkov V, Kholodenko BN. Reconstructing static and dynamic models of signaling pathways using Modular Response Analysis. Current Opinion in Systems Biology. 2018;9: 11–21. doi:10.1016/j.coisb.2018.02.003

10. Hu JX, Thomas CE, Brunak S. Network biology concepts in complex disease comorbidities. Nat Rev Genet. 2016;17: 615–629. doi:10.1038/nrg.2016.87

11. Huttlin EL, Bruckner RJ, Paulo JA, Cannon JR, Ting L, Baltier K, et al. Architecture of the human interactome defines protein communities and disease networks. Nature. 2017;545: 505–509. doi:10.1038/nature22366

12. Assi SA, Imperato MR, Coleman DJL, Pickin A, Potluri S, Ptasinska A, et al. Subtype-specific regulatory network rewiring in acute myeloid leukemia. Nat Genet. 2019;51: 151–162. doi:10.1038/s41588-018-0270-1

13. Pawson T, Warner N. Oncogenic re-wiring of cellular signaling pathways. Oncogene. 2007;26: 1268– 1275. doi:10.1038/sj.onc.1210255

14. Weinstein IB, Joe A. Oncogene addiction. Cancer Res. 2008;68: 3077–3080; discussion 3080. doi:10.1158/0008-5472.CAN-07-3293

15. Jimenez-Dominguez G, Ravel P, Jalaguier S, Cavaillès V, Colinge J. An R package for generic modular response analysis and its application to estrogen and retinoic acid receptor crosstalk. Sci Rep. 2021;11: 7272. doi:10.1038/s41598-021-86544-0

16. Klinger B, Sieber A, Fritsche-Guenther R, Witzel F, Berry L, Schumacher D, et al. Network quantification of EGFR signaling unveils potential for targeted combination therapy. Mol Syst Biol. 2013;9: 673. doi:10.1038/msb.2013.29

17. Gapp BV, Konopka T, Penz T, Dalal V, Bürckstümmer T, Bock C, et al. Parallel reverse genetic screening in mutant human cells using transcriptomics. Molecular Systems Biology. 2016;12: 879. doi:10.15252/msb.20166890

18. Subramanian A, Narayan R, Corsello SM, Peck DD, Natoli TE, Lu X, et al. A Next Generation Connectivity Map: L1000 Platform and the First 1,000,000 Profiles. Cell. 2017;171: 1437-1452.e17. doi:10.1016/j.cell.2017.10.049

19. Horvath S, Dong J. Geometric interpretation of gene coexpression network analysis. PLoS Comput Biol. 2008;4: e1000117. doi:10.1371/journal.pcbi.1000117

20. Obayashi T, Hayashi S, Shibaoka M, Saeki M, Ohta H, Kinoshita K. COXPRESdb: a database of coexpressed gene networks in mammals. Nucleic Acids Res. 2008;36: D77–82. doi:10.1093/nar/gkm840

21. Wang YXR, Huang H. Review on statistical methods for gene network reconstruction using expression data. J Theor Biol. 2014;362: 53–61. doi:10.1016/j.jtbi.2014.03.040

22. Faith JJ, Hayete B, Thaden JT, Mogno I, Wierzbowski J, Cottarel G, et al. Large-scale mapping and validation of Escherichia coli transcriptional regulation from a compendium of expression profiles. PLoS Biol. 2007;5: e8. doi:10.1371/journal.pbio.0050008

23. Meyer PE, Kontos K, Lafitte F, Bontempi G. Information-theoretic inference of large transcriptional regulatory networks. EURASIP J Bioinform Syst Biol. 2007; 79879. doi:10.1155/2007/79879

24. Margolin AA, Nemenman I, Basso K, Wiggins C, Stolovitzky G, Favera RD, et al. ARACNE: An Algorithm for the Reconstruction of Gene Regulatory Networks in a Mammalian Cellular Context. BMC Bioinformatics. 2006;7: S7. doi:10.1186/1471-2105-7-S1-S7

25. Meyer PE, Lafitte F, Bontempi G. minet: A R/Bioconductor Package for Inferring Large Transcriptional Networks Using Mutual Information. BMC Bioinformatics. 2008;9: 461. doi:10.1186/1471-2105-9-461

26. Butte AJ, Kohane IS. Mutual information relevance networks: functional genomic clustering using pairwise entropy measurements. Pac Symp Biocomput. 2000; 418–429. doi:10.1142/9789814447331_0040

27. Butte AJ, Tamayo P, Slonim D, Golub TR, Kohane IS. Discovering functional relationships between RNA expression and chemotherapeutic susceptibility using relevance networks. PNAS. 2000;97: 12182–12186.

28. Carette JE, Guimaraes CP, Varadarajan M, Park AS, Wuethrich I, Godarova A, et al. Haploid genetic screens in human cells identify host factors used by pathogens. Science. 2009;326: 1231–1235. doi:10.1126/science.1178955

29. Rane SG, Reddy EP. Janus kinases: components of multiple signaling pathways. Oncogene. 2000;19: 5662–5679. doi:10.1038/sj.onc.1203925

30. Bandyopadhyay S, Mehta M, Kuo D, Sung M-K, Chuang R, Jaehnig EJ, et al. Rewiring of genetic networks in response to DNA damage. Science. 2010;330: 1385–1389. doi:10.1126/science.1195618

31. Golub GH, Loan CFV. Matrix Computations. JHU Press; 2013.

